# Biofouling Characteristics of Graphene Oxide Membrane in a Protein-rich Environment

**DOI:** 10.1101/2021.05.30.446346

**Authors:** Richard Rode, Saeed Moghaddam

## Abstract

Membrane biofouling has inhibited permselective separation processes for decades, requiring frequent membrane backwash treatment or replacement to maintain efficacy. However, frequent treatment is not viable for devices with a continuous blood flow such as a wearable or implantable dialyzer. In this study, the biofouling characteristics of a highly hemocompatible graphene oxide (GO) membrane developed through a novel self-assembly process is studied in a protein-rich environment and compared with performance of a state-of-the-art commercial polymer membrane dialyzer. The studies are conducted in phosphate-buffered saline (PBS) environment using human serum albumin (HSA), which represents 60% of the blood protein, at the nominal blood protein concentration of 1 g L^-1^. Protein aggregation on the membrane surface is evaluated by monitoring the change in the membrane flux and SEM imaging. The GO membrane water flux declined only ~10% over a week-long test whereas the polymer membrane flux declined by 50% during the same period. The SEM images show that HSA primarily aggerates over the graphitic regions of nanoplatelets, away from the charged hydrophilic edges. This phenomenon leaves the open areas of the membrane formed between the nanoplatelets edges, through which the species pass, relatively intact. In contrast, HSA completely plugs the polymer membrane pores resulting in a steady decline in membrane permeability.

## 1. Introduction

Membrane biofouling has been the primary impediment to continuous operation of numerous separation processes.^[1–3]^ Formation of biofilms inhibits flux across the membrane, requiring increased operation pressure to maintain constant permeate flux. To mitigate biofouling issues and extend the membrane operation period, feed pretreatment,_[4,5]_ preventive overdosing of chemicals,^[6]^ and in-line backwashing, among other things, have been utilized. However, these strategies are not applicable in biomedical devices that interface with the blood flow continuously such as a wearable or implantable dialyzer.^[7–10]^ Hence, development of highly biofouling resistant membranes is required to maintain a steady hydraulic and sieving performance and prevent biofouling induced infections.^[11–13]^

Biofouling in biological media typically results from the formation of a thin biofilm consisting of various proteins amassed over the membrane surface. Polymer membranes exhibit extensive biofouling due to their high surface roughness^[14]^ and relatively hydrophobic characteristic,^[15–18]^ which encourages hydrophobic-hydrophobic interactions between the surface and proteins. Previous studies have indicated that this biofouling occurs over two defined steps. Initial protein aggregation occurs across the membrane surface and within the membrane pores, dictated by the membrane surface properties. This initial protein adsorption then acts as a nucleation site for further adsorption, dictated primarily by protein-protein interactions. Numerous studies have thus been dedicated to altering the membrane surface to prevent initial biofouling,^[19–24]^ as this is ultimately responsible for proliferation of protein adsorption. Alteration of the material surface has been pursued to reduce surface roughness to limit the cell and protein adhesion.^[25–28]^ Use of nanoparticles or bioactive materials has been explored for implantable devices or membranes.^[29,30]^ Current dialysis membranes employ a blend of hydrophobic-hydrophilic polymers to minimize biofouling. While the presence of these hydrophilic charged moieties has generally resulted in improved biofouling characteristic, recent studies have indicated that superhydrophobic surfaces can also extensively reduce protein adhesion.^[31,32]^ The extremely low surface energy of these materials is believed to deter protein adsorption by not meeting minimum interaction energies, leading to limited surface adsorption similar to hydrophilic structures.

Graphene oxide (GO) has been introduced in polymer mixed-matrix structures as a hydrophilic filler in an effort to reduce biofouling.^[33–35]^ These assemblies have demonstrated significant reduction in protein adhesion through minimization of available hydrophobic sites with expansion of hydrophilic surface moieties from the filler materials.^[36,37]^ Introduction of these hydrophilic sites across the hydrophobic surface improves surface wettability, limiting the available interaction sites for proteins through the presence of a thin water layer. These anionic sites mimic those typically found on the glomerular basement membrane of the kidney, which consist of >80% sulfated polyanions acting as permselective barriers to macromolecules in blood.^[38]^ The presence of these polyanionic sites indicates the effective screening of hydrophobic proteins via electrostatic rejection combined with limited available hydrophobic-hydrophobic interactions. The presence of hydrophilic filler materials also results in significant negative surface charge, further limiting protein adsorption through electrostatic screening of the negative charge typically found on protein molecules. While extensive research has been dedicated to investigating the impact of polymer surfaces on fouling, results concerning biofouling of stand-alone GO laminates over extended testing period and the mechanism governing the protein adsorption onto GO flakes and impacts on membrane flux have yet to be presented. Our previous work concerning this GO nanoplatelet assembly investigated its viability in blood-contact environments.^[39]^ Hemolysis, complement activation, and coagulation characteristics for the surface all fell within the physiological range, demonstrating the viability of the membrane in dialysis applications.

In this work, we investigate the biofouling characteristic of our trilayer GO membrane and compare its performance with a state-of-the-art commercial dialyzer. Oxidation extent across multiple GO surfaces is altered to assess impact on protein adhesion and the primary mechanism responsible for protein adsorption. Human serum albumin (HSA) is chosen as the protein foulant, as it constitutes 60% of the blood protein molecules concentration. HSA adsorption is found to be drastically lower in highly oxidized GO cases compared to commercial surfaces, attributed to the highly hydrophilic charged surface preventing hydrophobic-hydrophobic adsorption. These results are quantified via the flux recovery rate (FRR) and surface imaging for both MGO and dialyzer surfaces, indicating significant adsorption within the graphitic regions of MGO nanoplatelets. The limited decrease in FRR across testing is attributed to adsorption within the interior of the GO nanoplatelet away from the primary transport pathway, presumably due to the hydrophobic characteristic of the carbonaceous GO backbone structure.

## 2. Materials and Methods

Graphene oxide was synthesized from natural graphite flakes (Asbury^®^ Cat No. 3243) following Marcano’s method using 6:1 or 2:1 ratios of Graphite:KMnO_4_. Stable GO dispersion in a mixture of DI water (1 g L^-1^) was sonicated for 5.5 hours followed by centrifugation at 3000 rpm for 30 min. Poly(allylamine hydrochloride) (PAH, M_w_ = 17,500) was purchased from Aldrich and used as received. PAH (1 g L^-1^, pH = 4) was used as interlinker in these laminates due to the amine groups of the polymer readily binding to the carboxyl and hydroxyl functionalities present on GO surface.

## 3. Membrane Preparation and Fabrication of Biofouling Platform

Previous layer-by-layer (L-b-L) assembly of GO^[40–42]^ has been pursued in an attempt to minimize the membrane thickness and transport resistance. These L-b-L assemblies have achieved thicknesses of ~100 nm while maintaining permeabilities of ~25-60 mL hr^-1^ mmHg^-1^ m^-2^ with improved permselectivity of ionic molecules.^[43,44]^ However, these L-b-L assemblies have generally ignored the impact of GO nanoplatelet physicochemical properties on the transport characteristics of the membrane. From Figure 1A, an ideal GO membrane should consist of the fewest GO layers possible while minimizing surface defects which compromise membrane permselectivity.^[45]^ The transport pathway of the GO membrane is thus dictated by the nanoplatelet size, the distance between the GO layers, and the number of GO layers needed to eliminate defects across the surface. As shown in Figure 1A inset, smaller GO nanoplatelets are desired across the surface to limit the transport distance between flake edge, which act as the primary transport pathway within the assembly. The sizing of these GO nanoplatelets is then dictated by the support membrane pore size, requiring GO nanoplatelets which are slightly larger than the pores to avoid nanoplatelets entering the pore.

**Figure 1.**
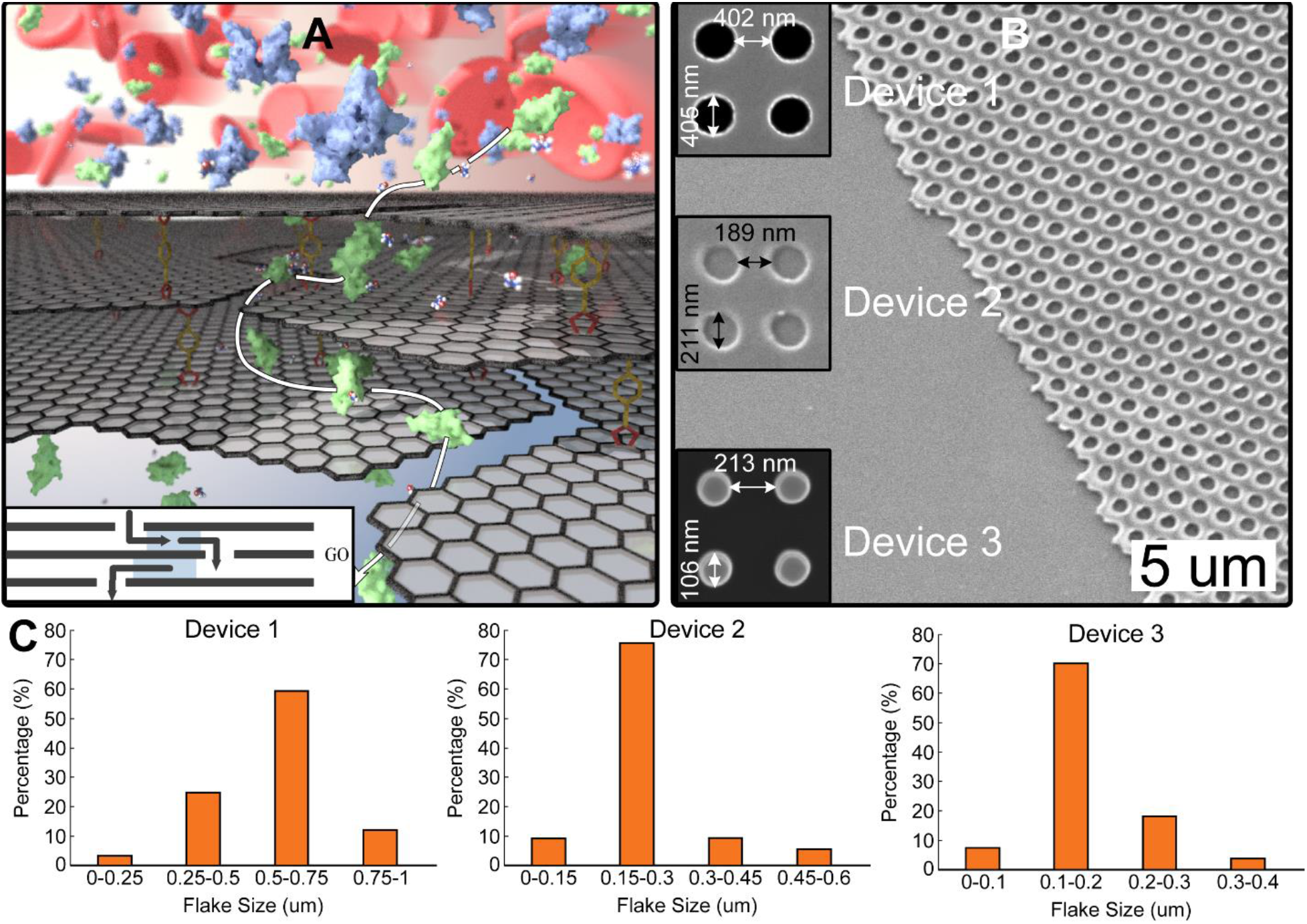
Visualization of GO layer-by-layer assembly with nanoimprinting of support membrane. (a) Transport pathway through layer-by-layer GO assembly, with inset showing impact of nanoplatelet size and interlayer spacing on transport. (b) SEM imaging of nanoimprinted pores on polymethyl methacrylate support membrane using various nano-feature sizes. (c) GO nanoplatelet size distribution for each tested device with varied pore size.

To allow for small nanoplatelet assemblies, ultra-thin (~200 nm) polymethylmethacrylate (PMMA) support membranes are utilized. 100 – 400 nm pores are introduced to the PMMA surface using nanoimprint lithography (NIL) stamps produced through electron-beam processing (Figure 1B). These pores are introduced into the PMMA membrane using an in-house imprinting system. The nanoimprinted PMMA membrane support is bonded to a polydimethylsiloxane (PDMS) film through oxygen plasma processing. Before bonding, the PDMS film introduces microchannels (50 μm width × 100 μm depth × 3 cm length) via polymer casting onto a previously machined polycarbonate master mold. For PMMA-PDMS bonding, the imprinted PMMA is initially exposed to oxygen plasma to introduce surface reactive moieties, then immediately immersed in 10% (v/v) (3-aminopropyl)triethoxysilane (3-APTES) solution for 15 minutes. The wafer is then removed from 3-APTES and again treated with oxygen plasma to allow bonding to PDMS under light pressure. The PMMA surface is hydrolyzed in 1.5M sodium hydroxide for 30 minutes at 45 °C to achieve the desired electronegative surface characteristic. With the support PMMA membrane processed, 120 – 500 nm GO nanoplatelets are produced through bath sonication for 5 – 10 hours (Figure 1C) to form the desired transport pathway demonstrated in Figure 1A inset.

While utilization of these small, highly sonicated GO nanoplatelets is key for producing ultra-permeable structures, the assembly process itself requires extensive optimization. Existing L-b-L assembly processes were initially tested to form an organized GO structure,^[46,47]^ but were ultimately unfit for sieving applications. These structures typically resulted in large open defect areas (dip-draw processing) or heavily stacked GO structures after a single immersion step (Figure 2A). As these techniques yielded poor sieving structures, many deposition steps would be needed to produce an efficient sieving surface at the expense of permeability. We have discovered within our research that utilization of small GO nanoplatelets (~100s nm) with sufficient diffusion time and processing conditions can produce ordered GO assemblies, yielding highly permselective membranes needing only three GO bilayers (Figure 2B). The self-assembly of these GO layers is dictated by the electrostatic interaction between individual nanoplatelets and the cationic interlinking molecule. GO nanoplatelets naturally tend to avoid overlapping due to the highly charged functionalities along their edges, resulting in electrostatic repulsion between nanoplatelets which minimizes that amount of overlapping while spacing nanoplatelets edge-to-edge dependent on their charge density. When the cationic interlinker (poly(allylamine hydrochloride) PAH) is introduced, electrostatic attraction between GO and PAH occurs, orienting the PAH normal to the charged surface, yielding an ~4 nm interlayer space (Figure 2C). By carefully controlling the pH, diffusion time of GO nanoplatelets, and concentration of PAH/GO solutions, the highly oriented GO structure observed in Figure 2B can be achieved. This process fundamentally differs from existing vacuum-filtration or dip-and-rise GO assemblies, as the GO nanoplatelet orientation associated with these processes are dictated by mechanical forces overcoming the relatively weak electrostatic forces between GO-GO or GO-PAH molecules. This process of introducing alternate solutions of GO and PAH given sufficient diffusion time to properly orient across the surface is carried out to produce the highly ordered GO structure observed. The hydrolyzed PMMA-PDMS device is initially immersed in PAH (1 g L^-1^) for 15 minutes to allow for proper orientation of cationic interlinker to the electronegative surface. Initial immersion times of 5- and 10-minutes were tested, but resulted in incomplete assembly atop the PMMA support, likely due to insufficient diffusion time of GO nanoplatelets that were weakly bonded to the surface. After 15 minutes, the device is removed and immersed in DI water in order to remove and weakly bonded PAH molecules. The device is then immersed in GO (1 g L^-1^) solution for an additional 15 minutes to allow sufficient diffusion and orientation of the sonicated GO nanoplatelets. The device is immersed once again in DI water to remove weakly bonded/misoriented GO nanoplatelets, where we define the presence of one GO-PAH layer as a single bilayer. This assembly process is repeated until three GO-PAH bilayers have been achieved atop the PMMA-PDMS device, resulting in limited defects or overlapping associated with previous layer-by-layer assembly processes. The previously fabricated PMMA-PDMS device is then alternately immersed in GO or PAH solutions for 15-minutes to allow for nanoplatelet organization, before being removed and washed with DI water to remove weakly bonded or misaligned nanoplatelets. Optimal immersion time was found to be 15 minutes, where initial tests of 5- and 10-minute immersions demonstrated noticeable defects across the membrane surface. Immersion for 15-minutes yielded the monolayer assemblyatop PMMA support with PAH interlinker. The optimal immersion time combined with significant washing following each GO stop results in the highly ordered nano mosaic observed in Figure 1B. Assembly of three GO-PAH bilayers atop the PMMA-PDMS device resulted in limited defect areas and overlapping associated with previous layer-by-layer assembly techniques.

**Figure 2.**
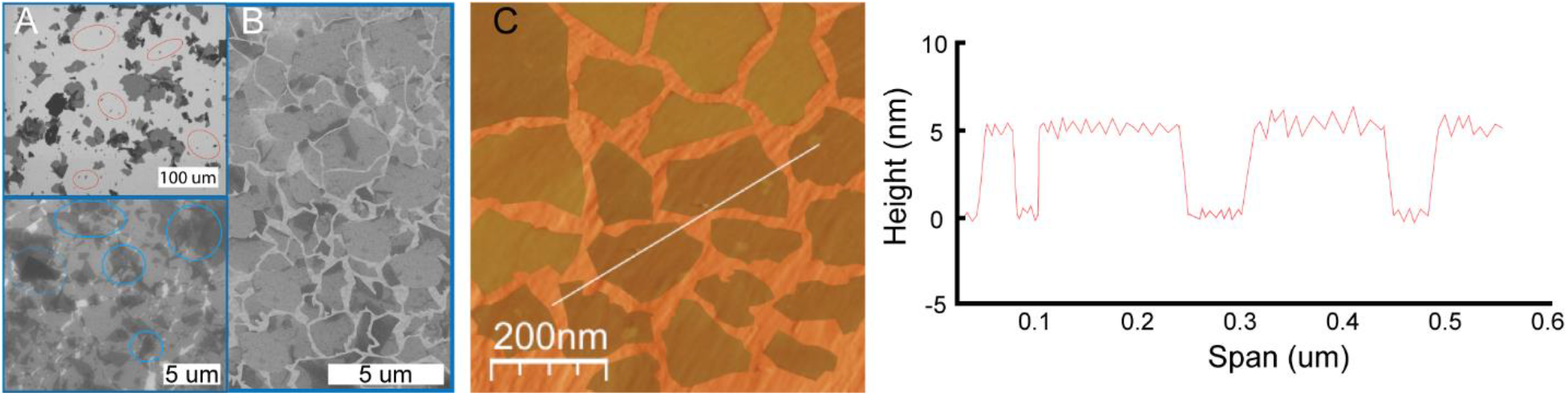
GO nanoplatelet assembly utilizing typical layering processes and varied immersion time. (a) Initial layering via dip-draw (top) or over-compacted structures (bottom). (b) Optimized assembly of GO nanoplatelets utilizing 15-minute immersion step. (c) AFM analysis of GO nanoplatelet anchored atop PMMA support using PAH interlinker.

The fully assembled GO device (Figure 3A-C) was evaluated for membrane permeability and selectivity of relevant dialysis species. Water permeability was determined for all three devices (Figure 3D), exhibiting improved permeability as GO nanoplatelet size was decreased up to a maximum permeability of 1562 mL hr^-1^ mmHg^-1^ m^-2^ for Device 3, nearly forty-fold greater than typical commercial high-flux dialysis membranes. To determine the permselective aspect of the GO device, urea, cytochrome-c, and Dextran T500 were employed as small- and middle-weight uremic toxins while retention of albumin was monitored (Figure 3E). B-galactosidase and phosphorylase-b were used to evaluate for the presence of large surface defects across the membrane, as their molecular weights are much greater than the molecular weight cut-off for the device itself. A maximum urea sieving coefficient of 0.5 was observed for Device 3 while all device iterations demonstrated >99% retention of human serum albumin (MW <66 kDa). Retention of β-galactosidase and phosphorylase-b were also >99.8% during testing, indicating that there were no significant surface defects or tears across the test devices (Figure 3F).

**Figure 3.**
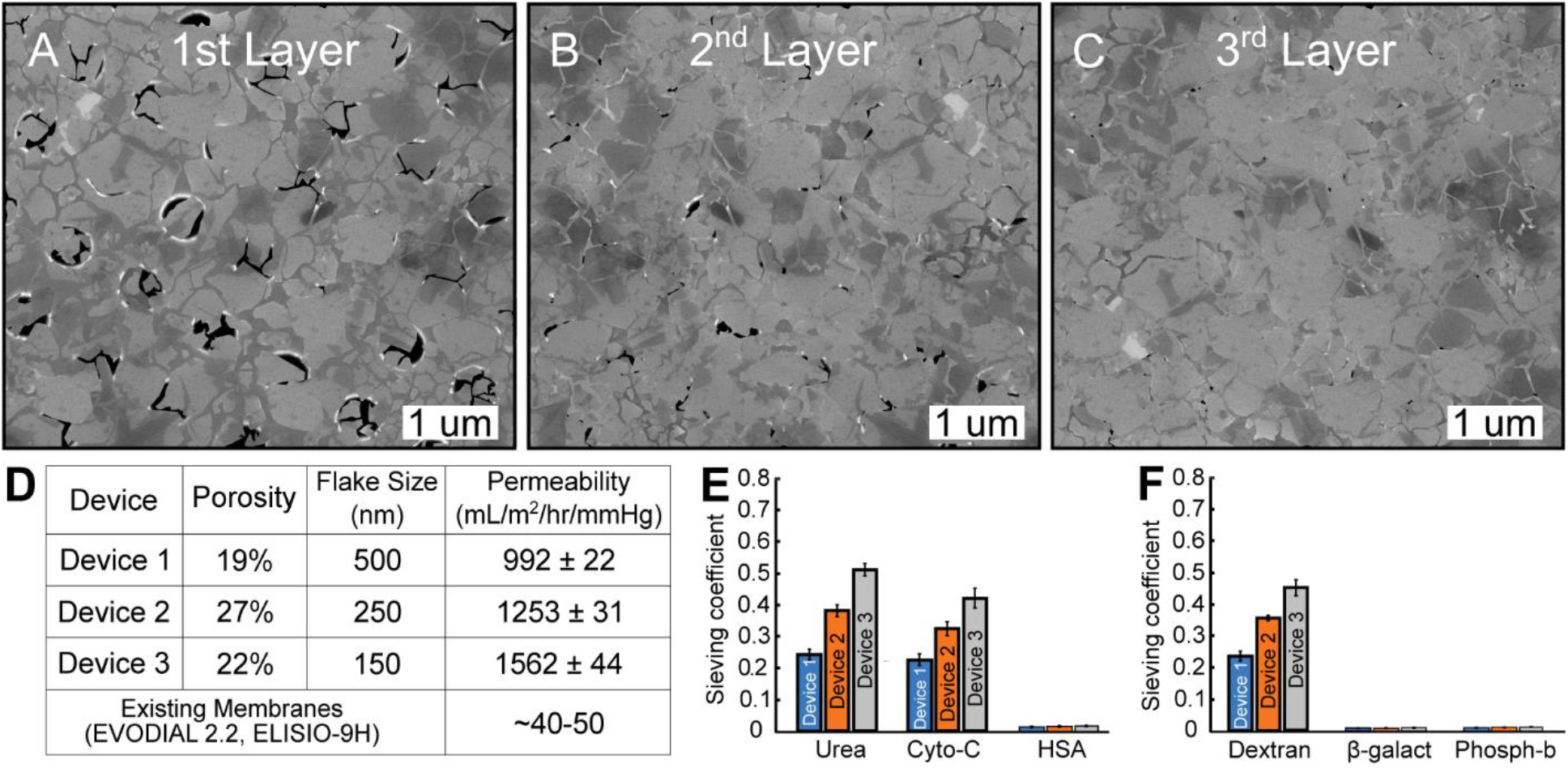
GO layer-by-layer assembly using three bilayers atop PMMA and permselective characterization of the membrane. (a-c) SEM imaging of subsequent layering of GO nanoplatelets atop PMMA support. (d) Device permeabilities and characteristics compared to commercial dialysis membranes. (e) Sieving coefficient for small- and middle-weight uremic toxins along with HSA loss. (f) Sieving characteristic of small-weight Dextran T500 and large-weight molecules, indicating no appreciable defects across the membrane surface.

The above fabricated microfluidic device was assessed for protein biofouling and flux recovery using an in-house setup. Pressure transducer data was monitored by data acquisition system (Agilent Technologies Inc., CA) at room temperature (T = 20 – 25 °C). Human serum albumin (HSA) solution (1 g L^-1^) was dissolved in phosphate-buffered saline (PBS) and supplied to the microfluidic device over a seven-day period. To monitor the impact of protein biofouling, DI water was run through the device for 10 min before monitoring the water flux of the membrane, recorded every six hours over the entire testing period. HSA concentrations were determined through UV-Vis spectroscopy at 280 nm after each step.

To determine the impact of HSA fouling on GO and polymer membrane structures, HSA dispersed in phosphate-buffered saline (1 g L^-1^) is passed through an in-house microfluidic test device over a seven-day period. The effective membrane area in each test was 9 mm^2^, with all tests carried out at room temperature (25°C) under 3 kPa supply pressure. To quantify the extent of protein fouling, flux for each membrane is evaluated every 6-hours, allowing a ten-minute span before monitoring flux to account for any shearing effects. The water flux before fouling (JBF) and throughout the protein fouling test (JAF) are recorded to estimate the flux recovery (FR%) of each membrane as follows:

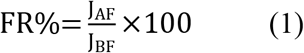

Biofouling mitigation typically relies on the surface roughness of the designated membrane alongside the surface charge/characteristic.^[48–50]^ A rougher membrane surface generally increases the tendency of fouling through clogging of the various valleys on the surface, inhibiting permeability of the membrane. AFM analysis for GO and polymer membranes demonstrates that the mean roughness (Ra) tends to increase from ~5 nm of GO to 12-15 nm of the polymer membranes. Water contact angle (WCA) and zeta potential measurements are used to determine the difference in oxidation state for the tested GO surfaces (Figure 4A, B). Both GO samples are produced following Marcano’s synthesis, where the difference in oxidation ratio used during the sample synthesis are denoted as MGO-1 (1:6 Graphite:Oxidizer) and MGO-2 (1:2 Graphite:Oxidizer). Figure 4A demonstrates the water contact angle of GO surfaces before exposure to protein fouling tests, demonstrating a highly hydrophilic surface typically associated with the oxidized GO surfaces. Figure 4B illustrates the zeta potential for both GO samples used, where typical mixed-matrix membranes employ highly hydrophilic (lower zeta) fillers to improve the surface wettability.^[51,52]^ MGO-1 demonstrates more electrostatically negative surface characteristic compared to the more sparsely oxidized MGO-2 surface, reinforcing the difference in surface charge of GO samples.

**Figure 4.**
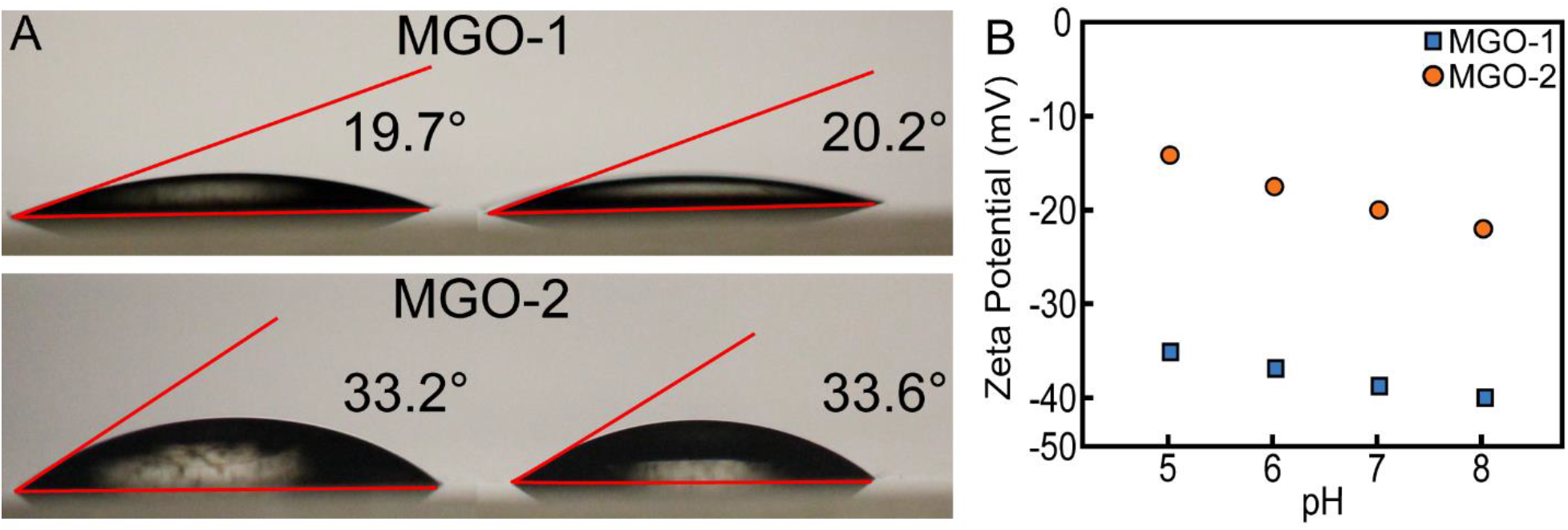
Surface characterization of MGO-1 and MGO-2 samples. (a) Water contact angle measurement of MGO-1 and MGO-2 used to visualize the surface wettability and (b) zeta potential of pristine MGO surfaces (evaluated on Paar Physica Electro Kinetic Analyzer).

## 4. Impact of Protein Fouling on Water Flux

In the previous work for this nano-assembly, initial blood-contact studies were carried out to quantify the impact of hemolytic behavior of GO-based nano mosaic membranes. Specifically, this work looked at the hemolysis, complement activation, and coagulation characteristics for various MGO surfaces while determining their viability in dialysis applications. The results observed in the previous work demonstrated MGO falls within the physiological range for all tested parameters, indicating a viable platform for use in dialysis applications. While the previous study provided valuable initial information concerning the hemocompatibility of MGO surfaces, evaluation of the biofouling characteristic is necessary to determine the efficacy of the membrane in protein-rich environments. Over the seven-day testing period, negligible decline in flux of the MGO-1 and MGO-2 membranes were observed compared to the polymer-based membrane examined. The flux for MGO-1 exhibited an ~10% reduction over the course of HSA testing compared to the ~14% reduction for MGO-2, while the polymer membranes demonstrated a flux of nearly 50% that of initial testing (Figure 5B). Significant decline in flux for both MGO-1 and MGO-2 surfaces is found within the first 24-hours of exposure to HSA, after which minimal change in flux is observed. Comparatively, commercial membranes demonstrate prolonged flux decline throughout the entire test interval, indicating continuous adhesion of HSA biofilms across the membrane. The nature of reduction in flux over the test period indicates an inherent difference in fouling mechanism between the GO and commercial membranes. This difference can likely be attributed to three main factors: 1) surface wettability, 2) surface roughness, and 3) membrane transport pathway structure. The GO membranes exhibit greatly improved surface wettability and surface roughness compared to the commercial membrane, inhibiting protein adhesion at the surface of the membrane.^[53–56]^ The surfaces of GO contain residual graphitic regions which remain unoxidized after synthesis, resulting in small hydrophobic regions within the inner graphitic sheet surrounded by oxygen-moieties at the flake edges (Figure 6A, B). This is in direct contrast to as-synthesized polymer membranes, consisting of hydrophobic surfaces which dehydrate the membrane surface and allow for significant protein adhesion.

**Figure 5.**
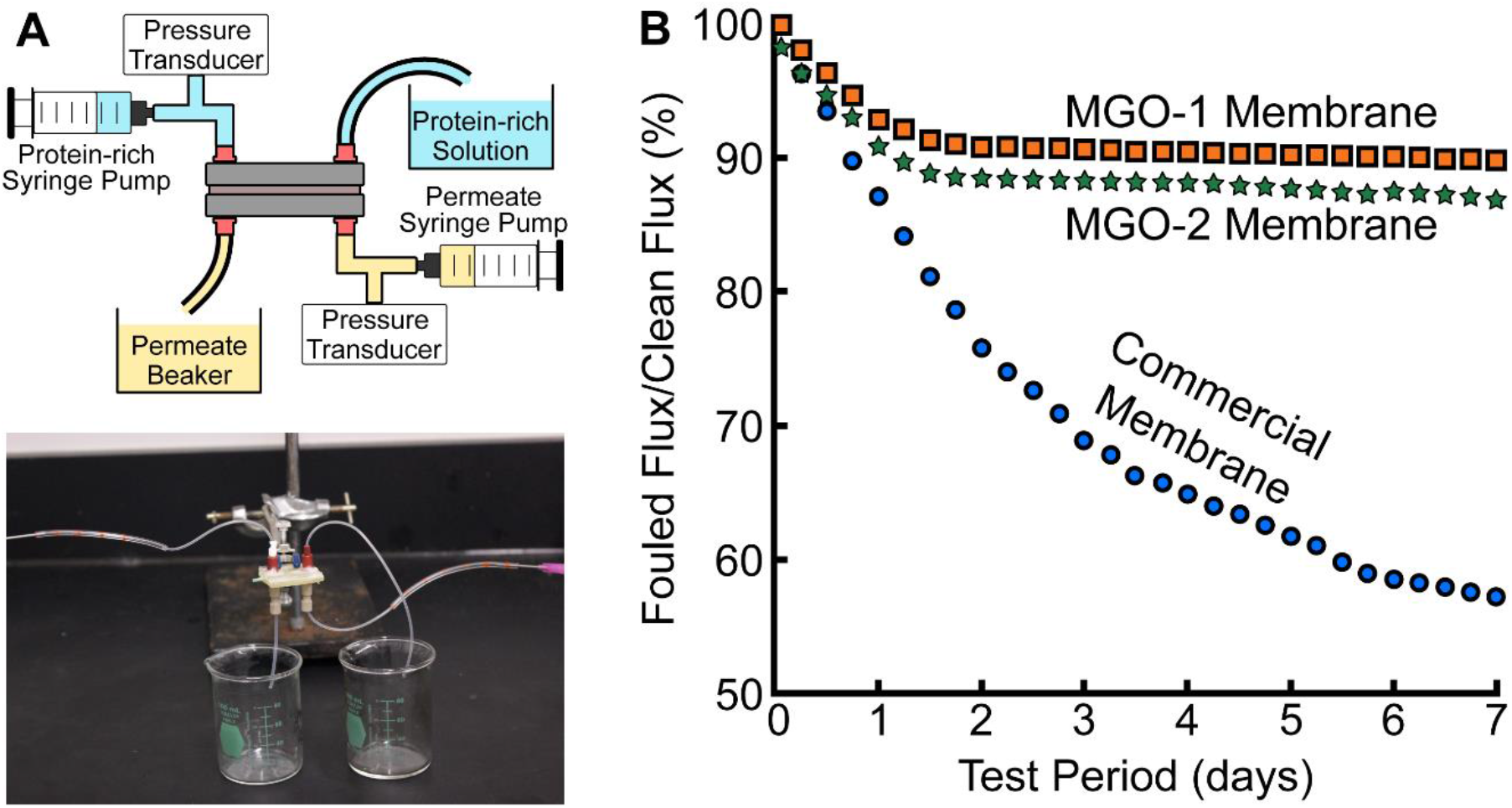
Fouling test schematic and membrane flux recovery during protein-rich testing. (a) Test device schematic and actual device photo. (b) Flux recovery for MGO-1, MGO-2, and commercial polymer membrane after 7-day testing exposure to 1 g L^-1^ HSA.

**Figure 6.**
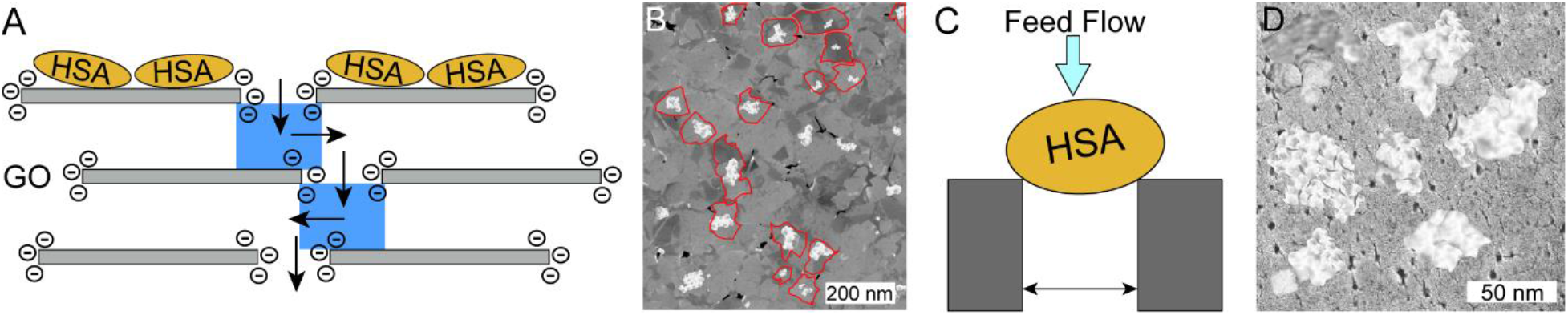
Schematic demonstration of protein adsorption across membrane surfaces. (a) HSA adsorption observed in GO assemblies, where primary adsorption occurs along the first GO layer within the graphitic flake regions away from hydrophilic edges. (b) SEM of HSA adsorption on GO layer with nanoplatelet edges highlighted. (c) Protein adsorption typical of polymer surfaces, showing proliferation of protein build-up across the surface and within pores. (d) HSA adsorption on commercial membrane surface after 7-day testing.

The flux trends observed for MGO-1 and MGO-2 can be attributed to the protein adhesion mechanism demonstrated in Figure 6A, where initial protein adhesion disrupts water transport through pore defects on GO surface introduced through synthesis.^[45,57]^ Permeation of these HSA proteins past the first GO screening layer are limited by the molecular weight cut-off inherent to the GO assembly, effectively preventing proliferation of adhesion within the bulk GO structure.

As the main transport mechanism within these GO layer-by-layer assemblies occurs at nanoplatelet edges and within the interlayer space,^[58]^ adhesion of HSA proteins within the hydrophobic graphitic backbone of the GO nanoplatelet results in relatively low impact on flux. In contrast, the commercial polymer membrane contains extensive hydrophobic sites for protein fouling, resulting in continuous protein adsorption and the steady decline in flux observed throughout the duration of testing (Figure 6C, D). Significant protein adhesion at the pore edge combined with secondary adsorption between protein molecules lends itself to a steady decline in the active pore radius of the membrane structured^[59–61]^ This effect of secondary adsorption occurs across the entire polymer surface, slowly degrading the primary transport pathway of the membrane, demonstrating the continuous decline in flux observed within this study. These adsorption phenomena are reflected in the SEM imaging of the tested membrane surfaces after exposure to HSA for the 7-day test period (Figure 7). Between the MGO-1 and MGO-2 scenarios, the impact of reduced oxidation extent is clearly visible within Figure 5B and 5C. The MGO-1 surface (Figure 7B) demonstrates limited HSA adhesion across the surface, which is limited to the hydrophobic flake regions. MGO-2 (Figure 7C) surface demonstrates significantly more adhesion compared to MGO-1, where HSA adhesion is found to bridge nanoplatelet edges, effectively blocking the primary transport pathway compared to MGO-1. While the protein adsorption within the graphitic regions across both samples are relatively similar, more adsorption near the flake edges is observed in the lesser-oxidized sample of MGO-2.

**Figure 7.**
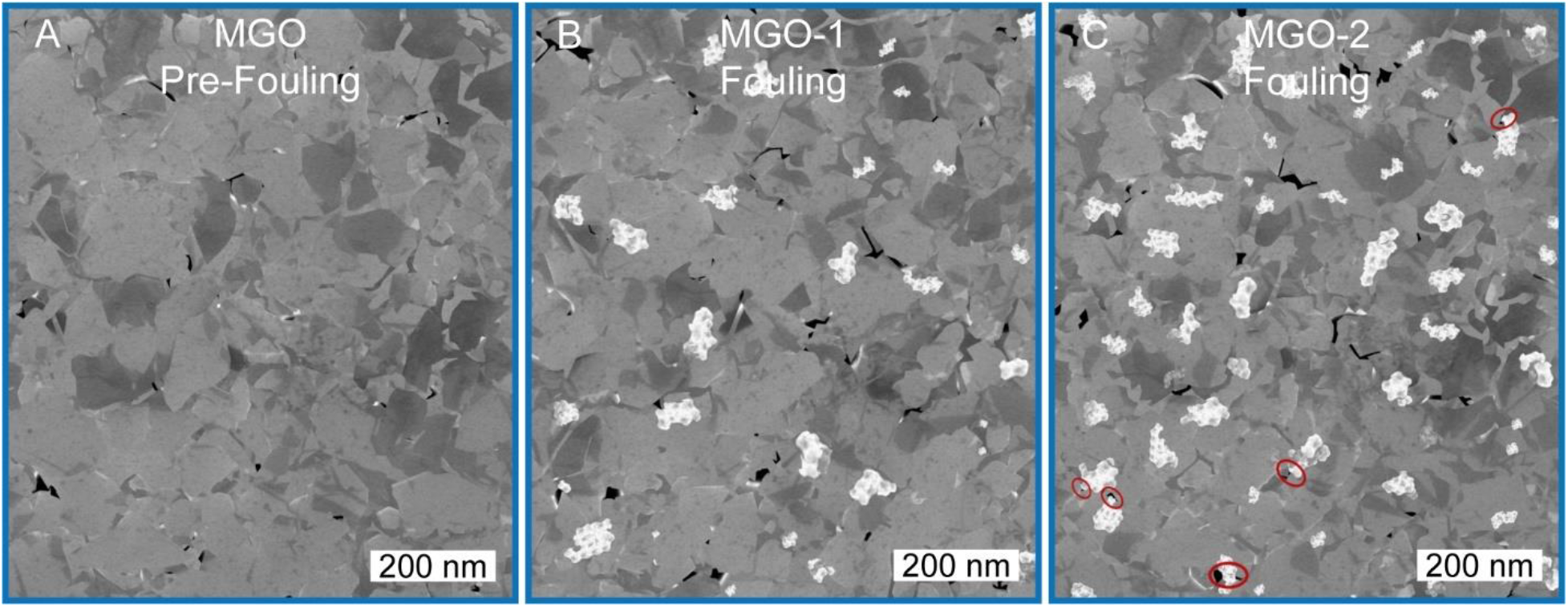
Visualization of HSA adsorption across tested membrane surfaces after 7-day exposure. (a) SEM image of MGO surface before exposure to HSA. Protein adhesion on (b) MGO-1 and (c) MGO-2 surfaces after exposure for seven-day test.

The lack of hydrophilic moieties at MGO-2 edges allows for slight adsorption of proteins near the flake edge, inhibiting the flux of the membrane over time. This adsorption characteristic provides insight into the flux trend observed for MGO-1 and MGO-2. The hydrophobic graphitic regions of the MGO samples are quickly occupied in the first 24 hours of exposure to HSA, resulting in the steep flux decline observed. Due to the difference in oxidation extent and lack of hydrophilic moieties at the nanoplatelet edges for MGO-2, a more significant decline in flux is observed. After the initial adhesion within these hydrophobic regions, flux recovery of the MGO membrane remains stable as the surfaces have become saturated with HSA, not allowing for further primary or secondary adhesion to occur within the subsequent GO layers. This is in stark contrast to typical polymer membranes, where primary and secondary protein adhesion can proliferate across the entire membrane surface and within membrane pores (Figure 6D), demonstrating the typical constant decline in flux recovery observed.

## 5. Conclusion

In summary, we have demonstrated the biofouling characteristic of a unique self-assembled GO nanoplatelet mosaic over a seven-day test period, providing insight into long-term protein adhesion characteristics of stand-alone GO membranes. These self-assembled GO mosaics demonstrate a decline in water flux of only 10%, vastly superior compared to the 50-60% loss typically attributed to commercial polymer membranes. This improved antifouling characteristic can be attributed to the presence of highly hydrophilic functionalities at GO nanoplatelet edges, inhibiting protein adsorption within the primary transport pathway of these GO laminates. The specific impact of these hydrophilic moieties is investigated through the adsorptive nature of two GO surfaces with varied oxidation extents. The MGO-1 sample exhibiting more extensive oxidation shows lower protein adsorption near flake edges compared to MGO-2, highlighting that the presence of proteins along the flake edge does inhibit the transport of water or other molecules through the membrane. The improved biofouling characteristic of these membranes provide a viable alternative to membranes exposed to protein-rich environments. This study demonstrates the efficacy of a surface-functionalized membrane in protein-rich scenarios compared to typical commercial membranes which blend these nanoparticles into the membrane matrix, retaining the permselective characteristic of the MGO surface.

## Supporting Information

Supporting Information is available from the Wiley Online Library or from the author.

## Acknowledgements

All authors have given approval to the final version of the manuscript. Research reported in this paper was supported by NIBIB of the National Institutes of Health under award number R21EB023527 to S.M. and T.R.G. The content is solely the responsibility of the authors and does not necessarily represent the official views of the National Institutes of Health.

S.M. was responsible for the project conception. R.P.R was responsible for all GO-based preparations, permeability testing, and protein adsorption experiments. R.P.R. wrote the manuscript and supplemental information with contributions from all authors. Authors declare no competing interests.

## Notes

### Competing Interest Statement

The authors have declared no competing interest.

